# DDR2 induces linear invadosome to promote angiogenesis in a fibrillar type I collagen context

**DOI:** 10.1101/2020.04.11.036764

**Authors:** Aya Abou Hammoud, Sébastien Marais, Nathalie Allain, Zakaria Ezzoukhry, Violaine Moreau, Frédéric Saltel

**Author notes:** Corresponding Author, INSERM U1053, Bariton, Université Bordeaux Segalen, 146 rue Léo Saignat, 33076 Bordeaux cedex, France. Laboratory of molecular Immunology and Signal Transduction, University of Liège, GIGA institute, CHU B34, 4000 Liège, Belgium.

## Abstract

To generate new vessels, endothelial cells (ECs) form invadosomes, which are actin-based microdomains with a proteolytic activity that degrade the basement membrane. We previously demonstrated that ECs form linear invadosomes in fibrillar type I collagen context. In this study, we aim to investigate the molecular mechanisms by which ECs guides angiogenesis in a fibrillar type I collagen context. We found that Discoidin Domain Receptor 2 (DDR2) is the collagen receptor tyrosine kinase required to form linear invadosomes in ECs. We further demonstrated that it acts in synergy with VEGF to promote extracellular matrix degradation. We highlighted the involvement of an interaction between DDR2 and the matrix metalloproteinase MMP14 in this process. Finally, using in vitro and *ex-vivo* angiogenesis assays, we demonstrated a pro-angiogenic function of DDR2 in a collagen-rich microenvironment. This study allows us to propose DDR2-dependent linear invadosomes as targets to modulate angiogenesis.

## Introduction

Angiogenesis is a process of new blood vessel formation from an existing vasculature. It is vital for growth, development, and healthy tissues maintenance. Angiogenesis is controlled by several cytokines such as vascular endothelial growth factor (VEGF), fibroblastic growth factor (FGF), epidermal growth factor (EGF) and platelet-derived growth factor (PDGF) that stimulate sprouting, proliferation and migration of endothelial cells (Rafii et al., 2002). To form new vessels, endothelial cells (ECs) have to degrade the extracellular matrix (ECM) in various contexts by forming structures called invadosomes (Daubon et al., 2016; Guegan et al., 2008; Seano et al., 2014a, 2014b). After degradation of the basement membrane, ECs have to progress in a different extracellular matrix composed by various components including fibrillar type I collagen.

Invadosomes are actin-based structures composed of actin regulatory proteins such as cortactin and N-WASP (neural Wiskott-Aldrich syndrome protein) and associated with scaffold proteins such as Tks5 (tyrosine kinase substrate 5) or RhoGTPases as Cdc42. Adhesion proteins such as integrins, talin or vinculin are also implicated in invadosome formation (Juin et al., 2014). ECs are able to form several types of invadosomes: dots, aggregates, rosettes (Seano et al., 2014a; Tatin et al., 2006) and linear invadosomes depending of the microenvironment (Di Martino et al., 2016; Ferrari et al., 2019; Juin et al., 2012; Schachtner et al., 2013). For example, invadosome rosettes in ECs are induced by VEGF or TGFß (Osiak et al., 2005; Seano et al., 2014a, 2014b; Tatin et al., 2006).

Invadosomes have the dual ability to interact with and degrade the ECM by recruiting matrix metalloproteinases (MMPs), such as MMP2, MMP9 and MT1-MMP (type 1 transmembrane protease, also known as MMP14) (Di Martino et al., 2016; Poincloux et al., 2009). MMP14 is a member of zinc-binding endopeptidases family. MMP14 presents a collagenase activity and can cleave type I, II and III collagens. It is expressed at the cell surface of various cell types, like cancer cells, endothelial cells and fibroblasts. MMP14 promotes cell migration, invasion, metastasis, angiogenesis and vasculogenesis, by degrading the pericellular ECM. MMP14 degrades ECM indirectly by activating other soluble MMPs including proMMP-2 and proMMP-13 (Haas et al., 1998; Han et al., 2016; Itoh, 2015; Lehti et al., 1998; Majkowska et al., 2017; Sabeh et al., 2004). Type I collagen in its fibrillar and physiological form behaves as a stimulus for MMP14 activation. Indeed, type I collagen fibrils stimulate MMP14 expression level, activation and localization along fibrils in several cell types including ECs (Di Martino et al., 2016; Juin et al., 2012; Majkowska et al., 2017). The mechanism by which type I collagen controls MMP14 in ECs is still not clearly understood. To support the relevance of our study, it is important to analyze ECs behavior in contact with fibrillar type I collagen and not with monomeric or degraded collagen like gelatin that can be use as control. Linear invadosomes are specifically formed upon contact to fibrillar type I collagen (Di Martino et al., 2016; Juin et al., 2014, 2012). In the context of type I collagen, Discoidin Domain Receptor 1 (DDR1) was involved in linear invadosome formation and activity in various tumor cells (Di Martino et al., 2016; Juin et al., 2014; Moreau and Saltel, 2015). But until now, there is no study reporting DDR2 role in invadosomes. Moreover, the collagen receptor involved in linear invadosome formation in ECs remains unknown.

Discoidin Domain Receptor 2 (DDR2) is among type I collagen-specific receptors that specially has an intrinsic tyrosine kinase domain. This receptor tyrosine kinase (RTK) is activated by autophosphorylation after interaction with type I collagen fibrils. It is mostly expressed in cells of mesenchymal origin such as fibroblasts and vascular smooth muscle cells (Vogel et al., 1997). DDR2, like its paralog DDR1, is considered as a collagen sensor and is involved in many biological processes in normal and cancer cells. DDR2 has an important role in proliferation, differentiation, adhesion, migration, invasion, metastasis formation and also in angiogenesis (Henriet et al., 2018; Hou et al., 2001). However, the role of DDR2 in angiogenesis is far from clear. Indeed, previous studies showed either a pro- or an anti-angiogenic effect of DDR2 (Badiola et al., 2012; Zhang et al., 2014; Zhu et al., 2015). Moreover, the mechanisms and cellular structures associated by which DDR2 guides angiogenesis should be pursued. It has been previously demonstrated that DDR2 overexpression increases *in vivo* tube formation (Zhang et al., 2014). Also, DDR2 mutant mice called slie (spontaneous autosomal-recessive mutation at Ddr2 locus, considered as DDR2 knockouts) lost the ability to form neovessels induced either by VEGF or tumor cells and this impairment is restored by the injection of DDR2-expressing adenovirus (Zhang et al., 2014). Furthermore, a strong reduction of pro-angiogenic (Vegfr2, Ang-2 and Mmp9) and a high increase of anti-angiogenic (Ang-1) candidates at mRNA levels were found in slie mice. Also, DDR2 negatively regulates the expression of pro-angiogenic factors (VEGF-A and Ang-2) at mRNA and protein levels. In contrast, DDR2 absence increased tumor angiogenesis in the liver as well as the expression of VEGF in hepatic stellate cells (HSC) derived from tumors (Badiola et al., 2012). In addition, DDR2 presents an anti-angiogenic effect in ocular disease (choroidal neovascularization-CNV). Indeed, an in vivo study by intravitreal injection of siDDR2 and DDR2-expressing adenovirus showed that DDR2 is responsible for the severity status of CNV (Zhu et al., 2015). Based on all these data, we suggest that the role of DDR2 in angiogenesis depends on cellular context driving the arrangement of ECs as tip and stalk cells which is the major key for angiogenesis initiation. Actually, the extracellular context is critical to determine which molecular mechanisms are activated during this arrangement. Here, we aim to investigate how DDR2 guides angiogenesis on fibrillar type I collagen context.

In this article, we reveal that DDR2 plays a decisive role in endothelial linear invadosome formation and activity in the presence of fibrillar type I collagen. In addition, we determine a molecular interaction between DDR2 and MMP14 directly involved in linear invadosome activity in ECs. Finally, we demonstrate that DDR2 silencing blocks angiogenesis in a type I collagen fibrils context. This study shows that DDR2 acts as a pro-angiogenic factor in a collagen-rich microenvironment through endothelial linear invadosomes and its interaction with MMP14.

## Results

### DDR2 expression and localization in endothelial cells

We previously demonstrated that ECs form linear invadosomes in a fibrillar type I collagen (Juin et al., 2012). Our objective was to determine if DDRs might be involved in their formation. First, we aimed to explore which DDR is expressed in endothelial cells. For that, DDR1 and DDR2 expressions were analyzed in different primary endothelial cell types (arterial, venous, lymphatic) and from different organs (umbilical vein, aorta artery, lung microvessels and dermis lymph vessels). In all ECs tested, in comparison to tumor A375 cells that express both DDR1 and DDR2, we found that a predominant expression of DDR2, especially in HUVECs (Figure 1A). We then addressed the ability of type I collagen fibrils to promote linear invadosome formation in ECs. Regardless of the source of ECs, we found that type I collagen fibrils strongly stimulate the formation of linear invadosomes (Figure 1B) and their ability to degrade ECM, as quantified using a classical *in situ* zymography assay. Indeed, the addition of type I collagen fibrils activated matrix degradation by ECs (Figure 1C).

**Figure 1:**
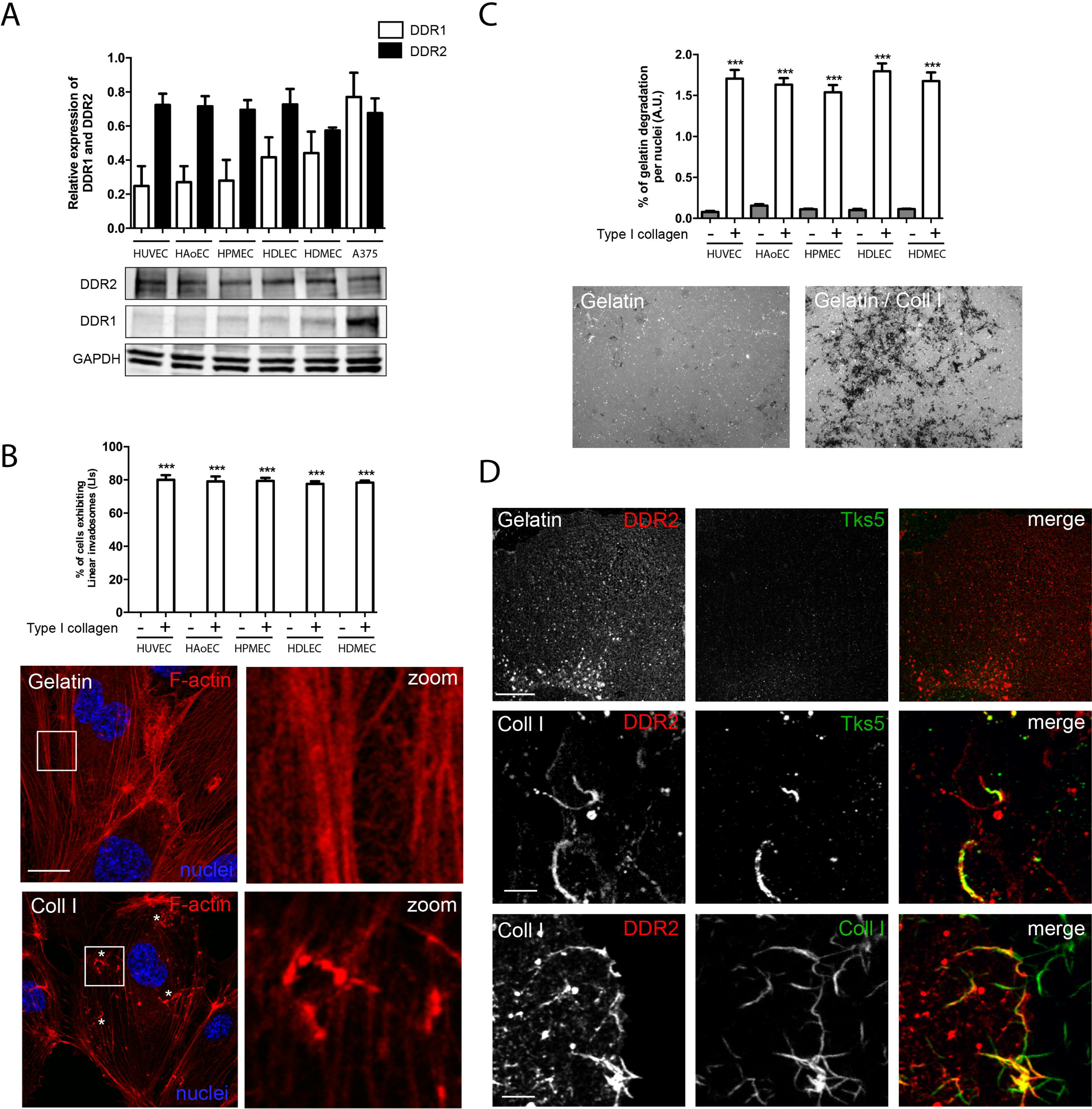
DDR2 expression and localization in endothelial cells. A) Comparaison of DDR1 and DDR2 expressions in endothelial cells. Cell lysates from HUVECs, HAoECs, HPMECs and HDLECs, HDMECs were analyzed by western blot. Tumor A375 cells were used as a positive control for DDR1 and DDR2 expressions. GADPH was used as a loading control. n = 3, Mean ± SD, ***P< 0.001. B) Fibrillar type I collagen promotes linear invadosome formation in all ECs. ECs were seeded on gelatin or on fibrillar type I collagen and the percentage of cells exhibiting linear invadosomes was quantified. n = 3, Mean ± SD, ***P< 0.001. The images represent HUVECs seeded on gelatin or on fibrillar type I collagen and stained with Phalloidin and Hoescht to reveal respectively actin (red) and nuclei (blue). Linear invadosomes are marked by an asterisk. The scale bar is 17 µm. C) Fibrillar type I collagen stimulates ECM degradation ability of all ECs. ECs were seeded on green fluorescent gelatin or on green fluorescent gelatin with fibrillar type I collagen and the ratio of gelatin degradation area per nuclei was quantified. n = 3, Mean ± SD, ***P< 0.001. The images represent the ECM degradation ability of HUVECs seeded on gelatin or on fibrillar type I collagen. Degradation areas are represented by the black zones. The scale bar is 50 µm. D) DDR2 localizes with linear invadosomes along type I collagen fibrils. HUVECs expressing DDR2 mcherry were seeded on gelatin (upper panel), on fibrillar type I collagen (middle panel) or on green fluorescent fibrillar type I collagen (lower panel). Cells were fixed and stained with anti-mcherry and anti-Tks5 (upper and middle panels) antibodies to reveal respectively DDR2 (red) and Tks5 (green). The scale bar is 4 µm.

HUVECs were used as cellular model for the whole study. Using immunofluorescence, we investigated DDR2 subcellular localization in HUVECs. Cells were seeded on gelatin or on fibrillar type I collagen. On gelatin, DDR2 exhibits a diffuse staining (Figure 1D upper panel), whereas DDR2 was localized along fibrils in type I collagen context (Figure 1D lower panel). As, type I collagen fibrils promote linear invadosome formation in ECs, we tested whether DDR2 colocalizes with the invadosome marker Tks5, and found that both proteins colocalize at these invasive structures in HUVECs. Similarly, Tks5 staining remains diffuse in HUVECS seeded on gelatin (Figure 1D middle panel). These data demonstrate that DDR2 is preferentially expressed in ECs and suggest its involvement in endothelial linear invadosomes induced along collagen fibrils.

### Type I collagen and VEGF enhance DDR2 expression and active invadosome formation

As VEGF plays a dual role in angiogenesis initiation, vessels branching and invadosome formation (Daubon et al., 2016; Guegan et al., 2008; Seano et al., 2014a, 2014b), we simultaneously tested the impact of VEGF treatment on DDR2 expression and on linear invadosome formation/activity. For this assay we established four cell conditions, in presence of gelatin (a denatured form of type I collagen, not recognized by DDR2) with or without VEGF or in the presence of type I collagen (on the top of a gelatin matrix) with or without VEGF (Di Martino et al., 2017). We first measured the impact of VEGF treatment on DDR2 expression (Figure 2A). We found that DDR2 expression is significantly and systematically increased upon VEGF addition independently of the ECM context. Type I collagen by itself enhances DDR2 expression. However, no synergic effect is noticed in type I collagen condition with VEGF. Interestingly, VEGF has no significant effect on DDR1 expression (Figure 2A). Thus, these results demonstrate that both VEGF and type I collagen promote DDR2 expression. Using the same conditions, we next investigated the impact of VEGF treatment on invadosome formation and activity. On gelatin, VEGF treatment promotes invadosome rosette formation in about 25 % of cells (Figure 2B), consistent with previous reports (Daubon et al., 2016). However, we found that DDR2 does not localize to these cortactin-positive structures (Figure 2B). In contrast, in collagen and VEGF condition, linear invadosomes were observed in more than 80% of cells (Figure 2B). Thus, if VEGF induces the invadosome rosette formation in gelatin condition, type I collagen is a potent inducer of linear invadosomes. In the collagen/VEGF condition, DDR2 still colocalizes with Tks5 on linear invadosomes (Figure 2B). Finally, using the *in situ* zymography assay, we measured and tested the impact of VEGF on invadosome degradation activity. The presence of type I collagen fibrils potentiates HUVECs degradation capacity, with a synergic effect in the presence of VEGF (Figure 2C). These data suggest a potential role of DDR2 in linear invadosome formation, in a type I collagen fibrils context.

**Figure 2:**
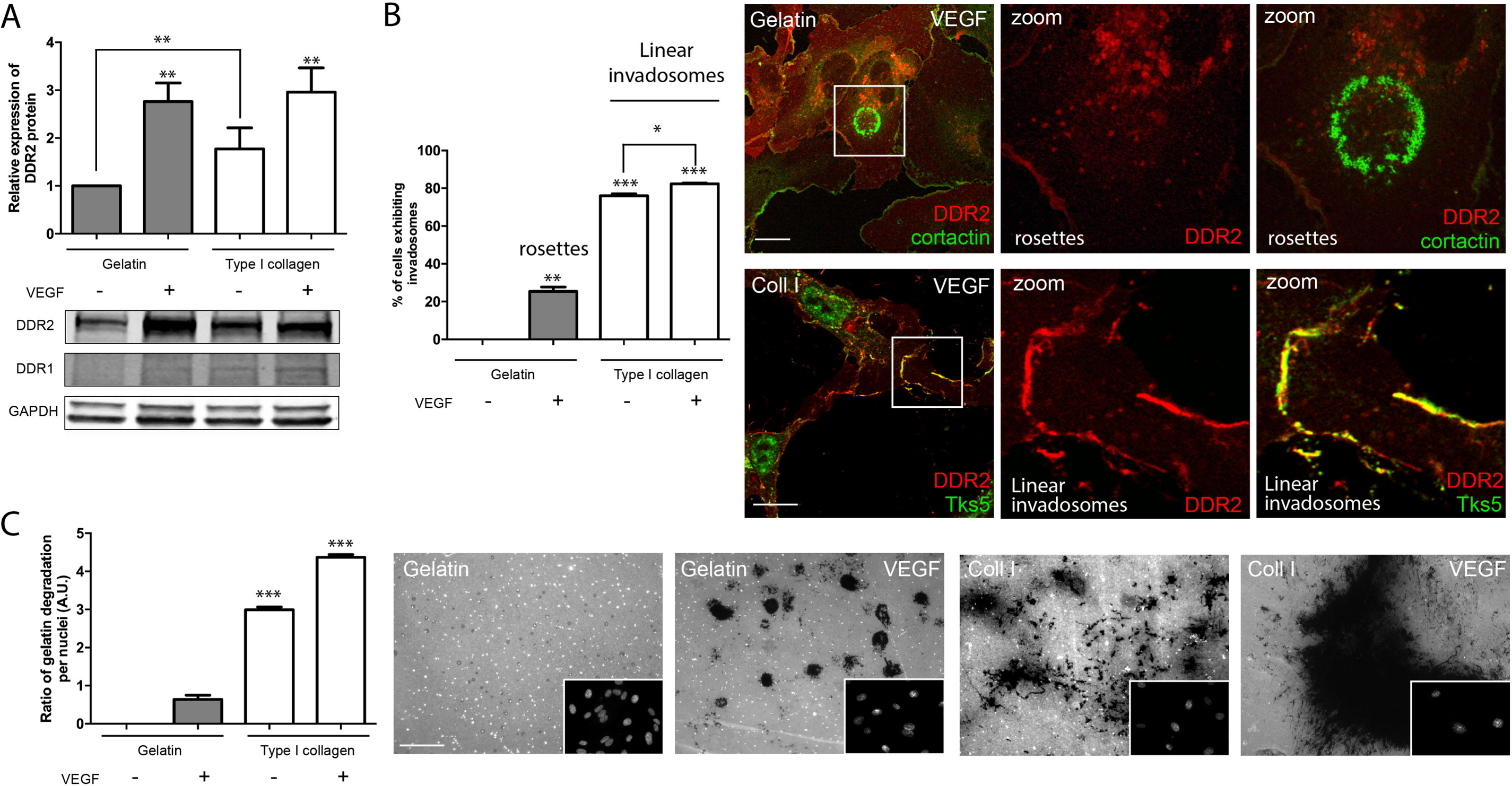
Effects of fibrillar type I collagen and VEGF on DDR2 expression and on invadosome formation/activity. HUVECs were seeded on two matrix types: gelatin or gelatin with fibrillar type I collagen and then treated or not 24 hours with VEGF (30 ng/ml). A) Comparaison of DDR1 and DDR2 expression in presence of fibrillar type I collagen and/or VEGF. Verification of DDR1 and DDR2 expression was performed by western blot of HUVEC lysats obtained from different conditions. GADPH was used as a loading control. n = 3, Mean ± SD, **P< 0.0012. B) Fibrillar type I collagen and VEGF enhance invadosome formation in HUVECs. Graph represents the quantification of cell percentage exhibiting rosettes or linear invadosomes. n = 3, Mean ± SD, *P< 0.0342; **P< 0.005; ***P< 0.001. Upon VEGF stimulation, DDR2 does not co-localize with rosettes on gelatin, but keep its co-localization with linear invadosome on fibrillar type I collagen. HUVECs expressing DDR2-mcherry were fixed and stained with anti-mcherry, anti-Cortactin and anti-Tks5 antibodies to reveal respectively DDR2 (red), Cortactin (green) and Tks5 (green). The scale bar is 21 µm. C) Fibrillar type I collagen and VEGF potentiate invadosome degradation activity in HUVECs. Degradation activity was analyzed by *in situ* zymography assay 24 hours after seeding. Graph represents the quantification of degradation activity ratio by counting gelatin degradation areas per nuclei of different conditions. n = 3, Mean ± SD, ***P< 0.001. The images show the degradation areas, presented by the black zones, with the nucleus stained by Hoescht corresponding of each field. The scale bar is 50 µm.

### DDR2 is required for active linear invadosome formation

In order to determine DDR2 involvement in linear invadosome formation and activity in ECs, DDR2 extinction was performed in HUVECs using two distinct siRNAs. DDR2 silencing was controlled and quantified by Western Blot (Figure 3A). Effective DDR2 depletion was associated with a decrease in the percentage of cells exhibiting linear invadosomes (Figure 3A). Whereas about 80% of cells exhibited linear invadosomes in the control condition (siCT), less than 30% were found in the presence of each siRNA targeting DDR2 (Figure 3A). The loss of linear invadosomes was associated with a degradation arrest using both siRNA against DDR2 (Figure 3B). We also demonstrate that in gelatin condition, DDR2 is not involved in rosette formation neither in their degradation activity in a VEGF-rich microenvironment (Figure S1). In addition, we show that VEGF treatment does not compensate the loss of DDR2 in the induction of active linear invadosomes in a type I collagen context (Figure S2). Altogether, these data demonstrate a specific role of DDR2 in linear invadosome formation and degradation activity exclusively in fibrillar type I collagen context.

**Figure 3:**
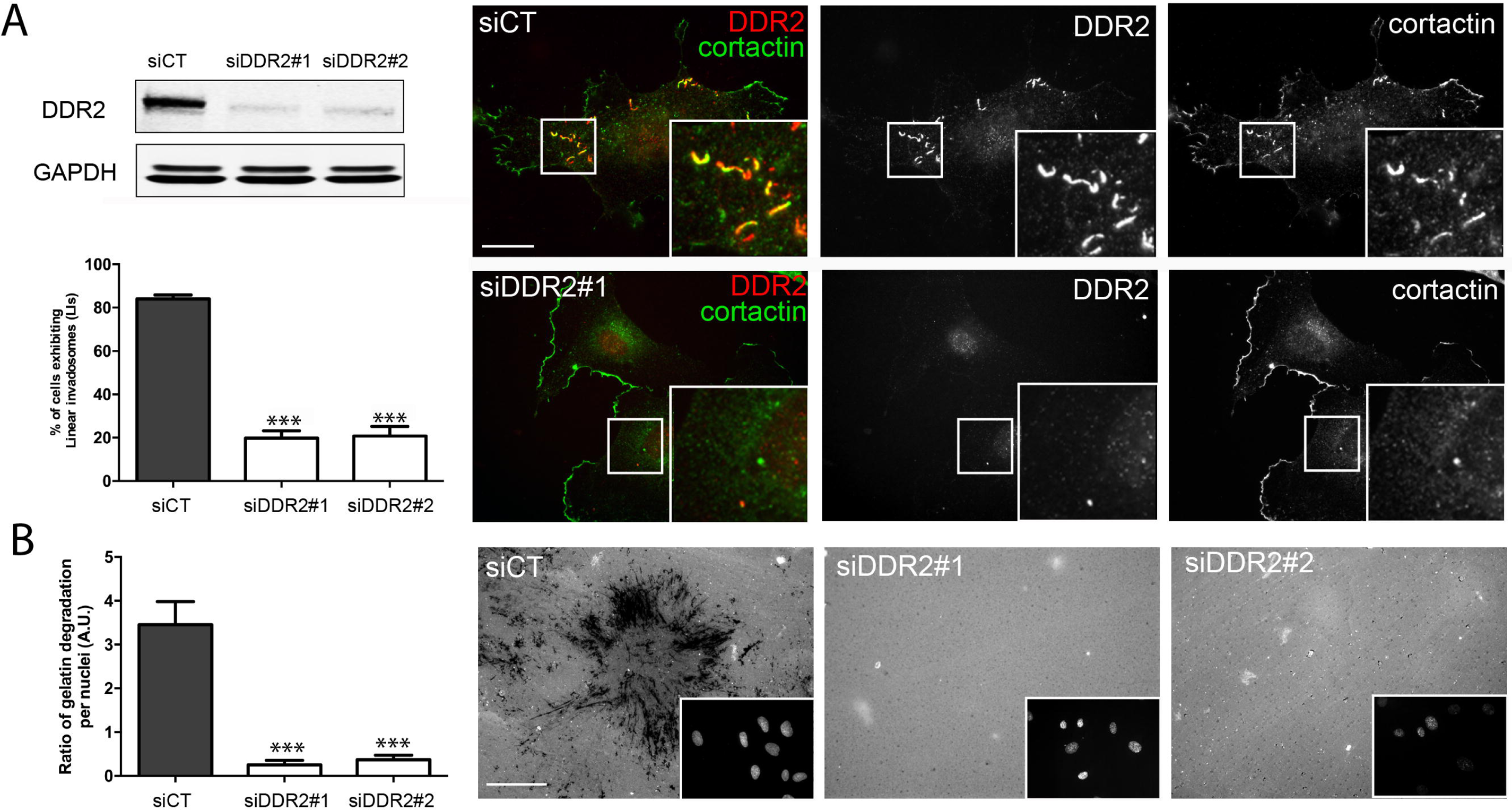
DDR2 extinction reduces linear invadosome formation and impairs HUVECs degradation activity in fibrillar type I collagen. HUVECs were transfected with three different siRNA: siCT as a control, siDDR2#1 and siDDR2#2 both targeting DDR2. A) After 48 hours, DDR2 silencing was validated by western blot. GADPH was used as a loading control. The percentage of cells exhibiting linear invadosomes was quantified. n = 5, Mean ± SD, ***p< 0.0001. Cells were fixed and stained with anti-DDR2 and anti-Cortactin, to reveal respectively DDR2 (red) and Cortactin (green). B) The degradation activity was analyzed by *in situ* zymography assay. The cells were seeded on fluorescent gelatin with fibrillar type I collagen matrix and observed after 24 hours. Degradation activity was quantified by counting gelatin degradation areas per nuclei. n = 3, Mean ± SD, ***p< 0.0009. The images show the degradation areas, corresponding to the black zones, with the nucleus stained by Hoescht, for the same field. The scale bars are 50 µm.

### DDR2 binds to MMP14 and regulates its activity in fibrillar type I collagen

Invadosomes have the ability to degrade ECM by recruiting MMPs, such as MMP9, MMP2 and MMP14. It has been demonstrated that type I collagen stimulates MMP14 expression and functions in several cell types including ECs (Haas et al., 1998; Itoh, 2015; Lehti et al., 1998; Majkowska et al., 2017; Sabeh et al., 2004). We thus investigated the link between linear invadosome-associated DDR2 and MMP14 in a fibrillar type I collagen context.

First, we validated by immunofluorescence that MMP14 colocalizes with DDR2 at linear invadosomes (Figure 4A). GM6001, an MMP inhibitor does not alter linear invadosome formation whereas it inhibits MMP14 recruitment and blocks their degradation activity (Supplemental Figure 3, A and B). Next, we tested if MMP14 and DDR2 do interact in ECs in the same context. By proximity ligation assay (PLA) we confirmed that DDR2 and MMP14 interact in fibrillar type I collagen. PLA green fluorescence is only observed when cells are incubated with an anti-mcherry targeting DDR2-mcherry and an anti-MMP14 antibodies comparing to the negative control or to the condition with each antibody separately (Figure 4B). Also, by co-immunoprecipitation assay, we showed that DDR2-mcherry and MMP14 co-immunoprecipitate (Figure 4C). MMP14 active forms at 55kDa and 28kDa are detected when DDR2-mcherry (about 130kDa) complex is immunoprecipitated and vice versa. Next, our objective was to determine whether MMP14 activity was mediated by DDR2 in ECs. MMP14 degrades ECM by activating other soluble MMPs including proMMP2 (Overall and Sodek, 1990; Sabeh et al., 2004). Using gelatin zymography assay, we evaluated MMP14 activity by detecting the latent (pro-MMP) and activated form of MMP2 (gelatinase A) and MMP9 (gelatinase B, MMP9 active form is known as a sign of MMP2 activation) (Lehti et al., 1998; Spinale, 2007; Stanton et al., 1998) upon DDR2 modulation. For that, DDR2 was either depleted or overexpressed by using, respectively, two distinct siRNAs or lentivirus-expressing DDR2-mcherry (Figure 4D and Supplemental Figure 3C). HT1080 fibrosarcoma cells treated with Concanavalin A (ConA) contain active MMP-2 (62 kDa) and served as a positive control (Toth et al., 2012). We showed that MMP2 and MMP9 are activated in fibrillar type I collagen as shown by the presence of their activated form at 62KDa and 85KDa respectively, but not when HUVECs are treated with GM6001 or cultured on gelatin (Figure 4D). Besides, we demonstrated that DDR2 extinction affects MMP2 and MMP9 activation whereas its overexpression increases the activation level of these two enzymes even in gelatin condition. These results were obtained in cell lysates as well as in supernatants (Figure S3C). These data show that DDR2 is required for collagen-induced MMP14 activation cascade. In parallel, we also checked MMP14 protein expression by western blot in the same conditions to reveal pro and active forms of MMP14. MMP14 is active in HUVECs and HUVECs DDR2-mcherry cultured on collagen or on gelatin by the accumulation of its active forms at 55kDa and 28kDa. In contrast, MMP14 is repressed upon GM6001 treatment (Figure S3C). In addition, we showed a decrease of MMP14 activated forms upon DDR2 extinction. 44kDa form of MMP14 is present in all conditions. This form behaves as a dominant negative regulator of the enzyme function and regulates the endocytosis of the active protease to preserve a viable level of MMP14 on the cell surface (Cho et al., 2008). All these results demonstrate that DDR2 is linked to and mediates MMP14 activity at linear invadosomes in ECs in a fibrillar type I collagen context.

**Figure 4:**
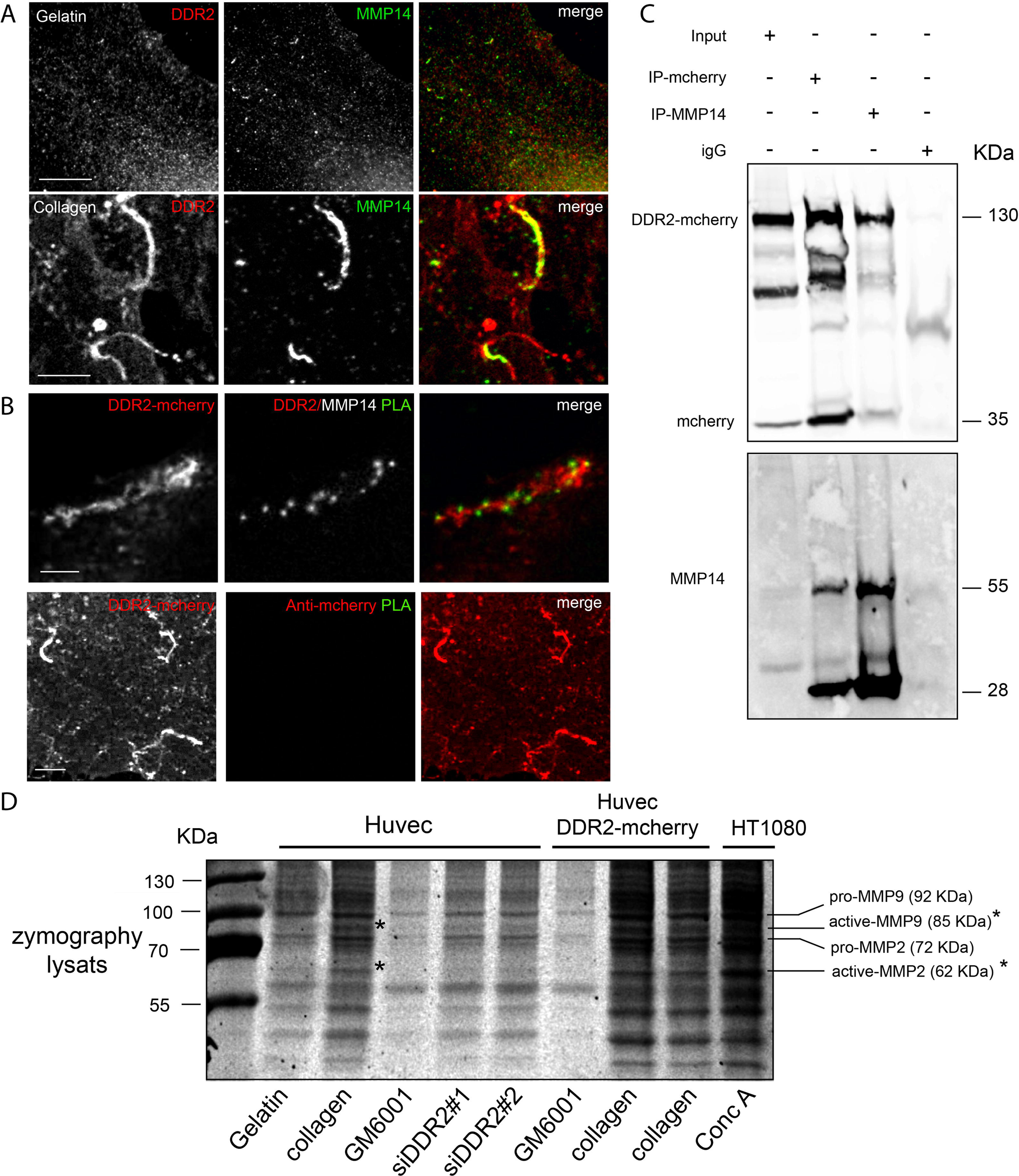
DDR2 activates MMP14 in fibrillar type I collagen. A) HUVECs were grown in the presence of fibrillar type I collagen for 24 hours and an immunofluorescence was carried out. Tks5 and MMP14 colocalization is marked with a primary anti-Tks5 and anti-MMP14 antibodies to reveal respectively Tks5 (red) and MMP14 (green). For MMP14 and DDR2 colocalization, DDR2 (red) and MMP14 (green) are marked respectively with a primary anti-DDR2 and anti-MMP14 antibodies. The scale bars are 50 µm. B) HUVEC-DDR2 mcherry cells were grown in the presence of fibrillar type I collagen for 24 hours and the proximity ligation assay (PLA) protocol was performed. The cells were stained with both anti-mcherry and anti-MMP14 primary antibodies to detect DDR2/MMP14 interaction (PLA green). For negative control, the cells were stained with anti-mcherry primary antibody alone. The scale bars are 2 and 5 µm, respectively. C) HUVEC-DDR2 mcherry cells were grown in the presence of fibrillar type I collagen for 24 hours. DDR2 and MMP14 were immunoprecipitated from protein extracts incubated with anti-mcherry and anti-MMP14 antibodies. The anti-IgG was used as a negative control. DDR2-mcherry and MMP14 were detected in whole lysats (Input) and in the immunoprecipitated proteins by western blot using anti-mcherry and anti-MMP14 primary antibodies. D) HUVECs were treated with two different siRNAs for DDR2 silencing or with DDR2-mcherry lentivirus for DDR2 overexpression. For MMP14 inhibition, HUVECs were treated with GM6001 (10 µM). HT1080 treated with Concanavalin A (ConA) serve as a positive control. All these cells were grown in the presence of gelatin or fibrillar type I collagen and in serum-free medium for 24 hours. The lysats were analyzed by gelatin zymography assay. Pro-MMP2, active MMP2 (asterisk), pro-MMP9 and active MMP9 (asterisk) were detected at 62kDa, 72 kDa, 85 kDa and 92 kDa respectively.

### DDR2 regulates angiogenesis *in vitro* and *ex vivo*, depending on MMP14

In order to analyze the role of DDR2-dependent linear invadosomes in angiogenesis, we used two models in parallel: HUVECs spheroids and *ex vivo* aortic ring assays (Figure 5 and 6). We tested the impact of DDR2 depletion or overexpression (DDR2-mcherry) on vessel formation. VEGF served as a positive control whereas siRNA targeting VEGFR2 (siKDR) and GM6001 inhibiting MMPs as negative controls. DDR2, MMP14 and VEGFR2 protein expressions were controlled by Western Blot in spheroids and in aortic rings (Figure 5A and 6A). New vessel formation in the presence of fibrillar type I collagen was then monitored by measuring vessel number and length.

**Figure 5:**
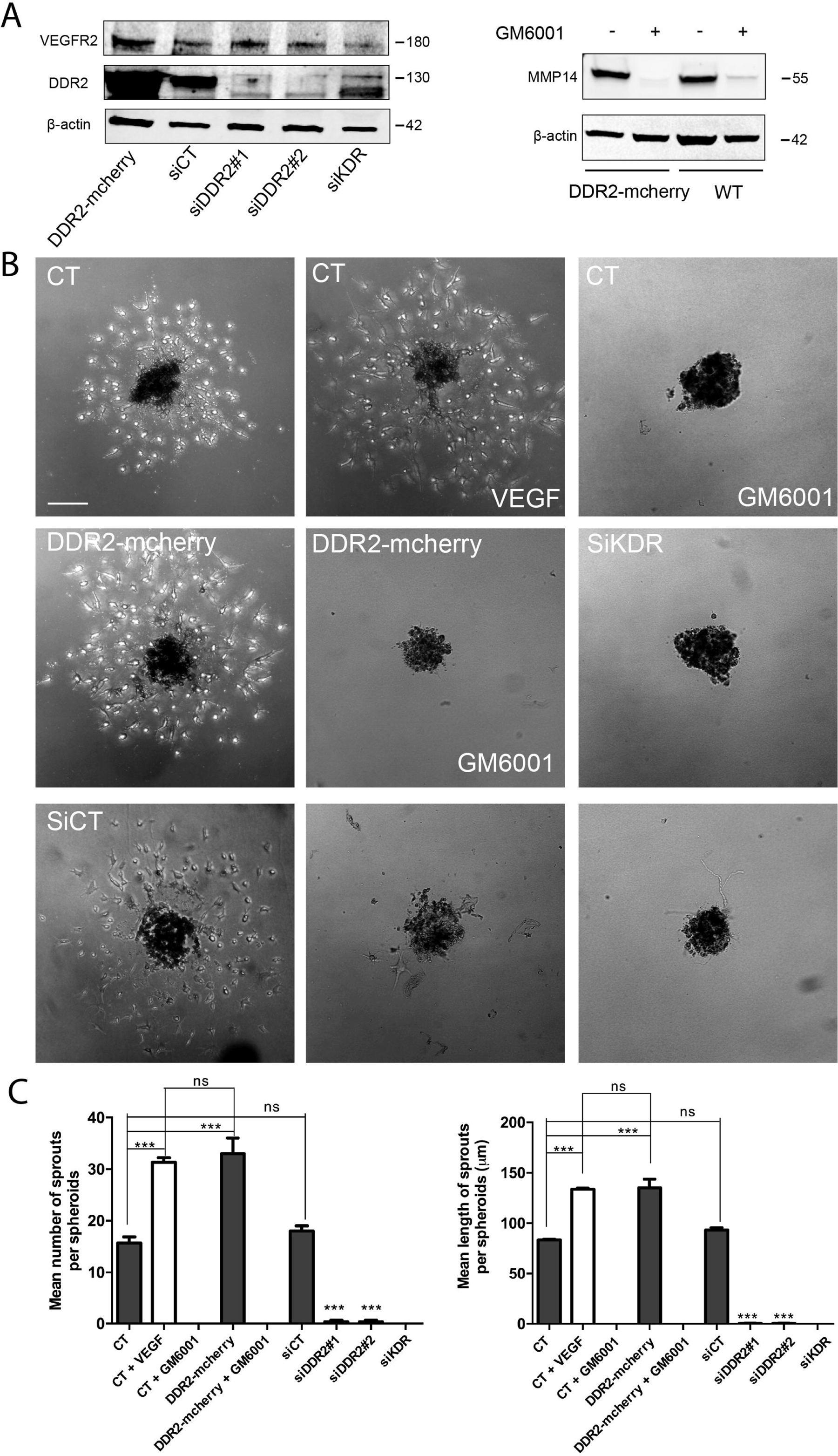
DDR2 and MMP14 regulate sprouting *in vitro*. HUVECs were transfected with siCT as a control, siDDR2#1 and siDDR2#2 for DDR2 knock-down and with siKDR for VEGFR2 silencing. For DDR2 overexpression, HUVECs were infected by DDR2-mcherry lentiviruses. For MMP14 inhibition, spheroids were treated with GM6001 (10 µM). A) After 48 hours, DDR2, VEGFR2 and MMP14 protein expressions were validated by western blot. β-actin was used as a loading control. One representative experiment out of three is shown. B) Spheroids obtained from HUVECs (siGL2, siDDR2, siKDR and DDR2-mcherry) were embedded in collagen (2 mg/ml) and treated with VEGF (30ng/ml) for the positive control. B) Sprouts outgrowth were visualized by observing EC spheroids in phase contrast. The scale bar is 21 µm. C) Quantification of total sprouts number and length was measured from at least 3 spheroids per experimental condition by using ImageJ software. Mean ± SD, ***P< 0.0001.

**Figure 6:**
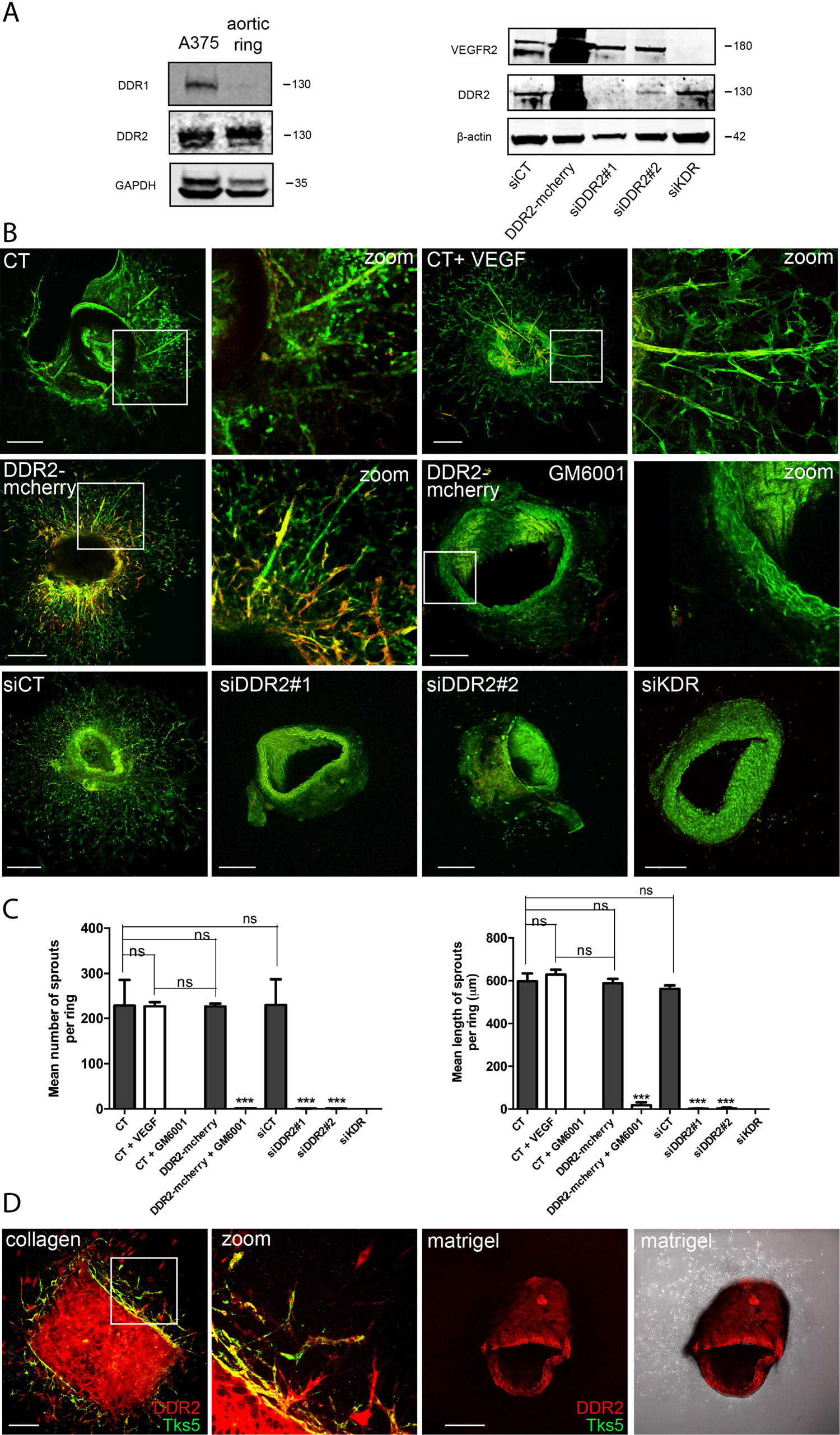
DDR2 and MMP14 induce neovessels formation *ex vivo*. A) Comparaison of DDR1 and DDR2 expressions in aortic rings. Protein lysates from mice aortic rings and A375 tumor cells (used as a positive control) were analyzed by western blot. GADPH was used as a loading control. One representative experiment out of three is shown. Aortic rings were transfected with control siRNA (siCT), DDR2 siRNAs (siDDR2#1 and siDDR2#2) or VEGFR2-targeting siRNA (siKDR). For DDR2 overexpression, the rings were infected by DDR2-mcherry lentiviruses. After 24 hours, DDR2 and VEGFR2 proteins expression were validated by western blot. β-Actin was used as a loading control. One representative experiment out of three is shown. B) Aortic rings were embedded in collagen (2 mg/ml). Rings treated with VEGF (30 ng/ml) serve as a positive control and those treated with GM6001 (10 µM) inhibit MMP14. Sprouts outgrowth were visualized by observing BS1 lectin (green) and nuclei (blue). C) Total sprout number and length were measured from at least 3 rings per experimental condition by using ImageJ software. Mean ± SD, ***P< 0.0001. D) For DDR2 colocalization with linear invadosomes, rings infected by DDR2-mcherry lentiviruses were used. Aortic rings were embedded in collagen (2 mg/ml) or in matrigel. DDR2 (Red) and Tks5 (green) were labelled with a primary antibody anti-mcherry and anti-Tks5 respectively. The scale bars are 21 µm.

In spheroids assays, in collagen condition, sprouts are formed. VEGF stimulation increases their mean number and length, whereas GM6001 blocks sprouts formation (Figure 5B and 5C). In parallel, DDR2 overexpression enhances sprouts formation with a mean number and mean length similar to those obtained with VEGF. Furthermore, MMP14 inhibition blocks sprouts formation also in DDR2-mcherry spheroids. At the opposite, DDR2 extinction suppresses new vessels formation as observed with negative control siKDR (Figure 5C and 5D). The results allow us to conclude that DDR2 and active MMP14 are both mandatory in the angiogenesis process in collagen context.

For the *ex vivo* ring aortic model, we first analyzed DDR1 and DDR2 expressions and confirmed that mouse ring aorta express predominantly DDR2 in comparison with A375 cells expressing both (Figure 6A). As for the spheroid assays, similar conditions were tested in the aortic model and we confirm in this model that collagen is able to promote sprouts formation in untreated aortic ring regardless of VEGF presence (Figure 6B and 6C). There is no significant difference in the mean number and mean length of sprouts per ring between control, DDR2 overexpression and VEGF addition (Figure 6B and 6C). Furthermore, we confirmed the induction of new vessels formation in the ring aortic with DDR2 overexpression. At the opposite, MMP inhibition, DDR2 and VEFGR depletion block this process (Figure 6B and 6C). These results are coherent to those observed *in vitro* with HUVECs spheroids. DDR2 major effect on vessels formation is restricted to fibrillar type I collagen. Interestingly, DDR2 and Tks5 labeling in aortic ring show a both expression of those markers in sprouts in collagen context. In matrigel condition DDR2 and Tks5 are not present in those structures (Figure 6D).

All these data prove that DDR2 and MMP14 regulate positively angiogenesis in fibrillar type I collagen/VEGF context. This essential role is mediated by endothelial linear invadosome formation and associated degradation activity.

## Discussion

DDR2 implication in angiogenesis is controversial and depends on cellular context. Besides, the mechanism by which DDR2 leads angiogenesis is still not clearly understood. Here, we report a pro-angiogenic function of DDR2 in fibrillar type I collagen context. We found that DDR2 positive activity in this process is related to its involvement in endothelial cell proteolytic activity linked to linear invadosome formation and associated matrix degradation. First, we established a specific DDR2 expression in a majority of endothelial cells. Using the HUVEC model, we validated that VEGF increases DDR2 expression independently of type I collagen presence. This is in accordance with a proteomic study that demonstrated an overexpression of DDR2 in HUVECs in response to VEGF stimulation (Katanasaka et al., 2007; Zhang et al., 2014). Besides, hypoxic conditions were shown to boost DDR2 expression at mRNA and protein levels (Zhang et al., 2014) which may explain the enhanced expression of DDR2 in tumor associated ECs during vascular initiation.

In ECs, we demonstrated that DDR2 localized with linear invadosomes induced by type I collagen fibrils. Also, we showed that combination of VEGF with fibrillar type I collagen enhanced linear invadosome formation and HUVECs degradation capacity. These data allow us to suggest that VEGF and fibrillar type I collagen contribute to increase DDR2 expression and activation promoting linear invadosome formation. In our study, we demonstrated that DDR2 is essential for active linear invadosome formation but not for invadosome rosette, in endothelial cells. These results confirm the matrix context specificity of this receptor to fibrillar type I collagen. Our data are in accordance with our previous results in cancer cells showing that DDR1 is localized with linear invadosomes and is essential for their formation and for their capacity to degrade ECM in the presence of fibrillar type I collagen, independently of its tyrosine kinase activity (Juin et al., 2014, 2012). Several studies have characterized the occurrence of invadosome rosette in endothelial cells in which their formation are influenced by different stimuli (Hai et al., 2002; Juin et al., 2013; Osiak et al., 2005). In HUVECs, invadosome rosette induced by VEGF or by Phorbol 12-myristate 13-acetate (PMA) depends on integrin α6β1 or on Src and Cdc42 activation respectively (Seano et al., 2014a; Tatin et al., 2006). Also, TGFβ long-term stimulation in BAE cells was able to stimulate rosette formation in which integrin αVβ3 plays a pivotal role (Varon et al., 2006). Recently, VEGF and type IV collagen were shown to stimulate rosette formation in human pulmonary microvascular endothelial cells (HMVECs) depending on Src and p190RhoGAP-B. These rosettes were involved in basal membrane degradation to initiate the angiogenesis process and also to connect vessels (Daubon et al., 2016; Juin et al., 2013; Seano et al., 2014a). Linear invadosomes could be implicated between these two steps to promote vessel progression by degrading the collagen helping tip cells to progress in stroma.

Our data indicate that collagen induces MMP14 activation through DDR2 in ECs model. We validated that MMP14 colocalizes with DDR2 at linear invadosomes and is essential for their activity but not required for their formation. Moreover, we demonstrated that DDR2 and MMP14 are linked and specifically in a type I collagen context. Also, we showed that MMP14 activity is regulated by DDR2 in this matrix environment. Indeed, we found that the presence of MMP2 and MMP9 active forms were DDR2 dependent. In addition, we showed a reduction of MMP14 activated forms upon DDR2 extinction. It has been reported that collagen-induced MMP14 activation is mediated through collagen-binding to integrins and DDRs, depending on cell types (Majkowska et al., 2017; Stanton et al., 1998). In stromal fibroblasts, DDR2 is a unique mechanism by which collagen-induced MMP14 activation, whereas, in cancer cells and epithelial cells, collagen upregulates MMP14 by other mechanisms including integrins (Ellerbroek et al., 1999). Previously, it has been demonstrated in human rheumatoid synovial fibroblasts (RASF) and human dermal fibroblasts (HDF) (both expressing only DDR2 but not DDR1) that MMP14 activation required only DDR2 signaling stimulation by fibrillar collagen because the selective DDR kinase inhibitor DDR1-IN-1 (a specific inhibitor used to block the kinase activity of DDR1 and DDR2) inhibits this activation (Majkowska et al., 2017). Also, it has been showed that Src-dependent phosphorylation of DDR2 at Tyr740 is important for DDR2 signaling upon collagen stimulation to induce MMP14 activation (Ellerbroek et al., 1999; Majkowska et al., 2017). Our data revealed that in ECs, DDR2 binds to and mediates MMP14 activity in fibrillar type I collagen context. This allowed us to suggest that DDR2 and MMP14 work as partners to regulate multiple functions including the formation of active linear invadosomes.

We demonstrated an important role of the couple DDR2-MMP14 during angiogenic sprouts formation in fibrillar type I collagen/VEGF contexts. With two distinct models, *in vitro* spheroids assays and *ex vivo* aortic ring assays, we showed that collagen induced sprouts formation in a DDR2 and MMP14-dependent manner. Sprouts formation was only observed in conditions whereby DDR2 and MMP14 are present and active. These data suggest that VEGF acts on endothelial cells in type I collagen context through DDR2 that is essential in angiogenesis. This DDR2 function related to linear invadosome formation and specially to their degradation activity is correlated with the colocalization with linear invadosome marker Tks5. This staining is only detected in aortic ring sprouts in collagen context. Despite all the studies that have demonstrated an important role of DDR2 in angiogenesis, our findings report for the first time the DDR2 pro-angiogenic effect dependent on the type I collagen context is due to linear invadosome formation. Moreover, it has been shown that α_6_β_1_ integrin calls up MMP14 at invadosome rosettes for vascular branching in tumor angiogenesis. Here, we indicated that DDR2 recruits MMP14 at linear invadosome to positively regulate sprouting formation demonstrating the plasticity of those structures. Recently, it was demonstrated that MMP14 pro-angiogenic impact is based on its capacity to interact with VEGFR1 which facilitate the binding of VEGFA to VEGFR2 to activate angiogenesis (Han et al., 2016). VEGF binds preferentially to VEGFR1 than VEGFR2, and thus, VEGFA stimulation of VEGFR2 needs a decrease in VEGFR1 to access on the cell surface. Also, in MMP14 knockout mice, MMP14 is pivotal for VEGFA-induced tube formation in vitro and corneal neovascularization in vivo. According to all these data, we hypothesize that fibrillar type I collagen activates DDR2 not only to stimulate MMP14 for endothelial proteolytic mechanisms, but, also to cluster and to activate MMP14 on specific sites at the cell surface.

Furthermore, the mechanisms by which DDR2 inhibits or activates angiogenesis remain poorly understood. Zhu et al. have showed a DDR2 anti-angiogenic effect in choroidal neovascularization-CNV due to its capacity to downregulate the phosphorylated level of PI3K/Akt/mTOR pathway (Zhu et al., 2015). Whereas, another team have demonstrated in pulmonary fibrosis that Angiotensin II-activated type I collagen expression in rat cardiac fibroblast is DDR2-dependent (George et al., 2016). In this context, they verified that Angiotensin II up-regulates DDR2 via NF-ΚB. Besides, this team have reported that DDR2 is critical for Angiotensin II or for TGFβ1 to promote lung vascular fibrosis by enhancing ERK1/2 MAPK phosphorylation (George et al., 2016; Ushakumary et al., 2019).

It is worth to note that one study demonstrated a link between DDR1 and angiogenesis in arterial wound repair. Hou et al. have demonstrated that SMCs isolated from DDR1-null mice impair the neointimal formation following arterial injury, *in vitro*. The authors explained this dramatic result that when DDR1 is absent, SMCs lose their capacity to attach to collagen and to activate MMP2 and MMP9 activities (Hou et al., 2001). Here, we hypothesize that in type I collagen context DDR1, as DDR2, regulate angiogenesis via linear invadosomes formation by recruiting MMP14.

In summary, all these data indicate a pro-angiogenic effect of DDR2 in collagen-rich microenvironment. We show that DDR2 regulating endothelial linear invadosome formation is determinant for sprouting angiogenesis. This allows us to propose DDR2 and linear invadosomes as new targets to modulate angiogenesis. Furthermore, our study highlights the benefits to determine the DDR2-dependent mechanisms which will be an important track to understand the microenvironment driven angiogenesis *in vivo* and consequently to improve targeted angiogenesis therapies.

## Materials and Methods

### Cell culture

HUVECs (Human Umbilical Vein Endothelial cells) provided by Laurent Muller (UMR7241 Centre interdisciplinaire de recherche en biologie, CIRB, Paris-France) were maintained in Endothelial Cell Gowth medium 2 with supplementMix (PromoCell, C22011). HAoECs (Human Aortic Endothelial Cells), HDLECs (adult Human Dermal Lymphatic Endothelial Cells), HPMECs (Human Pulmonary Microvascular Endothelial Cells) and HDMECs (adult Human Dermal Microvascular Endothelial Cells) were from PromoCell, Heidelberg, Germany (C-12271, C-12217, C-12281 and C-12212 respectively) and cultured in Endothelial Cell Gowth medium MV2 with supplementMix (PromoCell, C22022). For rosettes formation, HUVECs were treated with 30 ng/ml of VEGF (Recombinant Human Vascular Endothelial Cell Growth Factor, Fisher Scientific 10690920, Bordeaux, France) for 24 hours.

### Infection and transfection

For DDR2 localization, endothelial cells were infected twice in 2 days with lentiviral particles containing the plasmid PLVX-CMV-DDR2 mcherry at a multiplicity of infection of 5. For knock-down experiments, HUVECs or aorta were transfected with siGL2 (as a control), siDDR2-1, siDDR2-2 and KDR silencer® select (Ambion 4392420, Texas, United States). 30 μl of 20 μM siRNA were mixed with 30 μl of HiPerFect transfection reagent (Qiagen, Cat. No: 301704, Courtaboeuf, France) or Promofectine siRNA transfection reagent (PromoKine, PK-CT-2000-RNA-050, Heidelberg, Germany) and then incubated for 10 minutes at room temperature to allow the formation of transfection complexes.

### Invadosomes formation and degradation assays

Two types of matrix were produced under sterile conditions: gelatin and type I collagen fibrils for rosette and linear invadosome formation respectively. The coverslips were coated with gelatin (Sigma G1393) for 20 minutes at room temperature and fixed with 0.5% glutaraldehyde (Electron microscopy 15960) for 40 minutes at room temperature before being washed twice with PBS1X (Phosphate-Buffered Saline, GIBCO®). The type I collagen (Corning® Collagen I, rat tail cat. No: 354236, New York, United States) was diluted at 0.5 mg/ml in DPBS1X (Dulbecco’s Phosphate-Buffered Saline, GIBCO®) in order to add it to the gelatin. The whole was incubated for 4 hours at 37°C to allow collagen polymerization. Subsequently, the cells were inoculated at a concentration of 40,000 cells per well for 24 hours. For degradation activity test, we used the same protocol as above except that the coverslips were coated with fluorescent gelatin (Gelatin Dregon green life technologies G13186, Bordeaux, France).

### Western blot and Gelatin zymography assay

The cells or aorta were washed with ice-cold PBS and then lysed in radio-immunoprecipitation assay buffer (RIPA) composed of 0.1% SDS (Sodium Dodecyl Sulfate), 1% NP40, 150 mM NaCl, 1% sodium deoxycholate, 25 mM Tris HCl pH 7.4, and completed with anti-proteases (cOmplete™ ULTRA Tablets, Mini, EDTA-free, *EASYpack* Protease Inhibitor Cocktail, Roche®) and anti-phosphatases (PhosphoSTOP, Roche®). After incubation for 30 min at 4°C, extracts were centrifuged at 13,000 g for 15 min at 4°C and then the supernatant containing the proteins was recovered. For aorta protein extraction, tissues were crushed with a 1.5 ml tube pestle before centrifuge. The samples were assayed by the LOWRY method, prepared at 40 μg in Laemmli 4X buffer (8% SDS, 40% glycerol, 20% β-mercaptoethanol, 0.04% bromophenol blue, Tris HCl (pH 6.8) 500mM), heated at 95°C for 5 min, and loaded onto 10% SDS-PAGE gel (TGX™ Fast Cast™ Acrylamide Kit 10%, BioRad, California, United States). Proteins were transferred onto a nitrocellulose membrane using the Trans-Blot® Turbo™ semi-liquid rapid transfer system (BioRad), according to the supplier’s recommendations, blocked with Odyssey buffer (Blocking Buffer, LI-COR® Biosciences) for 1h, and probed with the primary antibody overnight at 4°C. The primary antibodies were diluted in TBS1X-0.1%Tween + 5% BSA: 1/1000 for DDR2 (Cell Signaling 12133S), phospho-DDR2 (R and D systems MAB25382, Minneapolis, United States) and DDR1 (Cell Signaling 5583S, WZ Leiden, Netherlands), FLK-1 (sc-6251, Santa Cruz, Texas, United States), MMP14 (Abcam 51074, Cambridge, United Kingdom) and 1/3000 for GAPDH (sc-166545, Santa Cruz) and β-actin (sc-47778, Santa Cruz).

Membranes were washed with TBS1X-0.1%Tween, incubated 2h at room temperature with the corresponding secondary antibody coupled to a fluorophore (IRDye® LI-COR®, Nebraska, United States) diluted to 1/5000, and then revealed with Odyssey® system (LI-COR Biosciences). Protein signals were quantified by ImageJ software.

For zymography assay, the protocol was adapted from Toth et al. (Toth et al., 2012). The cell lysates and the supernatant were loaded on SDS-PAGE gel (10% polyacrylamide/0.1% w/v gelatin) after preparing them with 4X non-reducing sample buffer. The gels were incubated 30 min with renaturing solution, were washed 2X with distilled water, were rinsed 30 min with developing buffer at room temperature with agitation and were incubated O/N with fresh developing buffer at 37°C in a close tray.

The gels were then stained with staining solution for 1h at room temperature with agitation, washed two times with distilled water and finally destained with destaining solution until bands can clearly be seen (Areas of enzyme activity appear as white bands against a dark blue background).

### Immunofluorescence

This protocol was performed as described in Di Martino et al. (Di Martino et al., 2017). The primary antibodies diluted 1/100 in PBS1X-4%BSA: DDR2 (Cell Signaling 12133S), Tks5 (fish M-300 sc30122, Santa cruz), Cortactin (clone 4F11 Millipore 5180) and MMP14 (Abcam 51074 and sc-373908). The cells were imaged using the Leica SP5 confocal microscope (63x/NA 1.5 plan Neofluor objective lens) and Carl Zeiss AxioCam MRc microscope. The images were analyzed by ImageJ software. Invadosomes formation and degradation assay are described in the supplemental Appendix.

### Co-immunoprecipitation

For MMP14 immunoprecipitation, HUVEC-DDR2 mcherry were used. The lysates were obtained from the cells cultured in collagen matrix at 2 mg/ml. After two washes with cold PBS1X, lysis buffer was added and the cells were scraped and incubated for 30 minutes on ice. The lysates were sonicated and centrifuged for 15 minutes. The proteins were loaded at 3 mg in each condition and incubated with the anti-IgG (negative control) and anti-MMP14 antibodies overnight at 4°C on the wheel. Next, the magnetic beads were added for 1 hour. After, immunoprecipitated protein-beads were washed 5 times with completed RIPA. The immunoprecipitated proteins were then recuperated with Laemmli 4X. Input was prepared using 50 μg of whole protein lysates. Proteins were incubated for 5 minutes at 95°C before electrophoresis.

For DDR2 immunoprecipitation, we used RFP-Trap_MA kit (rtmak20RFPTrap®_MA Kit (20 reactions), lot no.: 70404002MAK) from Chromotek, Planegg, Germany and the immunoprecipitation were done according to the manufacturer’s protocol.

### Proximity ligation assay

For Proximity ligation assay, we used Sigma-Aldrich kit from Merck-Sigma, Saint Quentin Fallavier, France. The experiments were done according to the manufacturer’s protocol. The primary antibodies were added after dilution at 1/50 with Duolink^®^ Antibody Diluent: DDR2 (Cell Signaling 12133S), Mcherry (NBP1-96752SS from Novus) and MMP14 (Abcam 51074, sc-373908).

### Spheroid sprouting assay

HUVECs were suspended in adequate culture medium with 20% methylcellulose (M7140 Sigma-Aldrich), added by drops (800 cells per 20 µl) and incubated overnight to form a single spheroid. Spheroids were collected, seeded in ECGM basal medium supplemented with 2 mg/ml of rat tail collagen, 20% FBS, 0.7% methylcellulose, NaOH 0.1 M to turn the mixture basic, 10 mM HEPES, 7.5% sodium bicarbonate and then incubated at 37°C and 5% CO_2_ for 1hour. After collagen polymerization, the complete medium was added to the samples. Spheroids treated with VEGF (30 ng/ml) were used as positive controls. Sprout outgrowths were imaged 48 hours later with a Leica SP8 laser scanning confocal microscope. Total sprout length was measured from at least 3 spheroids per experimental condition on phase-contrast images by using ImageJ FIJI software.

### Mouse aortic ring assay

This protocol was adapted from Baker M et al. (Baker et al., 2011) article. The aorta rings were cut at 0.5 mm and transferred in 5 ml of Opti-MEM without serum in a 10 cm dish. For transfection, the rings were incubated overnight at 37°C and 5% CO2 in Opti-MEM without serum and without penicillin/streptomycin.

For aortic ring embedding, the collagen was diluted on ice at final concentration of 2 mg/ml in DMEM 1X and the pH was adjusted with 20 μl per 10 ml of NaOH 5N to turn the mixture basic. 250 μl of collagen matrix or matrigel were transferred to each well of a 24-well plate. The plate was left for 10 min at room temperature and then incubated at 37°C and 5% CO2 for 1h. The embedded rings were then carefully feeded with 150 μl of Opti-MEM culture medium supplemented with 2.5% FBS and penicillin-streptomycin. Rings treated with VEGF (30 ng/ml) were used as positive controls. The growth medium was changed every 2 days by removing 130 μl of old medium and replace with 150 μl of fresh medium for 9 days.

For aortic ring immunostaining, the whole plate was washed with PBS containing CaCl_2_ and MgCl_2_. The aortic ring was fixed with 4% paraformaldehyde for 20 min at room temperature and then was permeabilized with PBS containing CaCl_2_, MgCl_2_ and 0.25% Triton X-100 for 15 min at room temperature. Blocking was performed for 30 min at 37°C in PBS1X containing 0.25% casein, and 0.015 M sodium azide. Primary antibodies were added and incubated overnight at 4 °C. We used 0.1 mg/ml of BS1 lectin-FITC to stain endothelial cells, 1/1000 anti-actin α-smooth muscle and 1/1000 of anti-DDR2 and anti-Tks5 antibodies. For primary antibodies uncoupled to fluorophore, the secondary antibodies were added and incubated for 3h at room temperature. DAPI was used for nucleus staining. After each antibody incubation, the ring was washed three times with PBS containing 0.1% Triton X-100 for 15 min and with distilled water for the last wash. Images were acquired with a Leica SP8 laser scanning confocal microscope. The images were analyzed by ImageJ FIJI software.

### Statistical tests

Data reported as the mean ± SEM of at least 3 experiments. Statistical significance (P< 0.05 or less) was determined using a paired *t-test* and analysis of variance (two-way ANOVA) as appropriate and performed with GraphPad Prism software.

## Acknowledgements

This research was funded by La Fondation pour la Recherche Médicale (grant number DEQ20180839586) by the Institut National du Cancer (INCA, grant PLBIO2015-140) and by Integrated Cancer Research (SIRIC, BRIO). We thank Bordeaux Imaging Center (BIC) for the support to this study.

## Author contributions

Aya Abou Hammoud performed all the experiments of this article, prepared the figures, wrote and edited the manuscript. Sebastien Marais contributed to images acquisition for in-vitro and ex-vivo angiogenesis assays. Nathalie Allain prepared the ring aortas from mice for ex-vivo angiogenesis assay. Zakaria Ezzoukhry performed preliminary experiments at the basis of the presented work. Violaine Moreau reviewed and edited the manuscript. Frédéric Saltel designed the project, analyzed the data, prepared the figures and wrote the manuscript.

## Conflicts of Interest

The authors declare no competing financial interests.

## Expanded View Figure Legends

**Figure S1:** DDR2 extinction does not impair rosette formation and activity. HUVECs were transfected with control siRNA (siCT) or siRNA targeting DDR2 (siDDR2#1 and siDDR2#2) and treated 24 hours with VEGF (30 ng/ml). A) After 48 hours, DDR2 extinction was validated by western blot. GADPH was used as a loading control. Rosette formation was assayed by immunofluorescence. Cells were fixed and stained with Phalloïdin and anti-Tks5 to reveal respectively Actin (red) and Tks5 (green). Rosette formation was quantified according to the percentage of cells exhibiting invadosome rosettes. n = 3, Mean ± SD, ns = not significant. B) The degradation activity was analyzed by *in situ* zymography assay. The cells were seeded on fluorescent gelatin matrix and then were observed after 24 hours. Degradation activity was quantified by counting gelatin degradation areas per nuclei. n = 3, Mean ± SD, ns = not significant. The images show the degradation areas, presented by the black zones, with the nucleus stained by Hoescht corresponding of each field. The scale bars are 50 µm.

**Figure S2:** VEGF does not restore the DDR2-mediated impairment of active linear invadosomes. HUVECs were transfected with control siRNA (siCT) or siRNA targeting DDR2 and treated 24 hours with VEGF (30 ng/ml). A) After 48 hours, DDR2 extinction was validated by western blot. Linear invadosome formation was assayed by immunofluorescence. Cells were fixed and stained with Phalloïdin and anti-Tks5 to reveal respectively Actin (red) and Tks5 (green). The percentage of cells exhibiting linear invadosomes was quantified. n = 3, Mean ± SD, ***p < 0.0001. B) The degradation activity was analyzed by *in situ* zymography assay. The cells were seeded on fluorescent gelatin and fibrillar type I collagen matrix and then observed after 24 hours. Degradation was quantified by counting gelatin degradation areas per nuclei. n = 3, Mean ± SD, ***p < 0.0001. The images show the degradation areas presented by the black zones with the nucleus marked by Hoescht corresponding of each field. The scale bars are 50 µm.

**Figure S3:** DDR2 and MMP14 interaction is required for active linear invadosome formation in fibrillar type I collagen. A) HUVECs were treated with GM6001 (10 µM) and were grown in the presence of fibrillar type I collagen for 24 hours and an immunofluorescence was carried out. Tks5 and MMP14 colocalization is marked with a primary anti-Tks5 and anti-MMP14 antibodies to reveal respectively Tks5 (green) and MMP14 (red). For MMP14 and DDR2 colocalization, DDR2 (green) and MMP14 (red) are marked respectively with a primary anti-DDR2 and anti-MMP14 antibodies. The scale bar is 50 µm. B) The degradation activity was analyzed by *in situ* zymography. The cells treated with GM6001 were seeded on fluorescent gelatin and fibrillar type I collagen matrix and observed after 24 hours. The images show the degradation areas, presented by the black zones, with the nucleus stained by Hoescht corresponding of each field. The scale bar is 50 µm. C) HUVECs were treated with two different siRNAs for DDR2 silencing or with DDR2-mcherry lentiviruses for DDR2 overexpression. For MMP14 inhibition, HUVECs were treated with GM6001. HT1080 treated with Concanavalin A (ConA) serve as a positive control. All these cells were grown in the presence of fibrillar type I collagen and in serum-free medium for 24 hours. The supernatants were analyzed by gelatin zymography assay. Pro-MMP2, active MMP2 (asterisk), pro-MMP9 and active MMP9 (asterisk) were detected. Protein expressions of MMP14 active form 55kDa and 28kDa were analyzed and were quantified by western blot from protein extracts. β-Actin was used as a loading control. n = 3, Mean ± SD, ***p < 0.0001.

## Notes

### Competing Interest Statement

The authors have declared no competing interest.

