## Supplementary figures and images for "DDR2 induces linear invadosome to promote angiogenesis in a fibrillar type I collagen context"

### fig sup 1.jpg

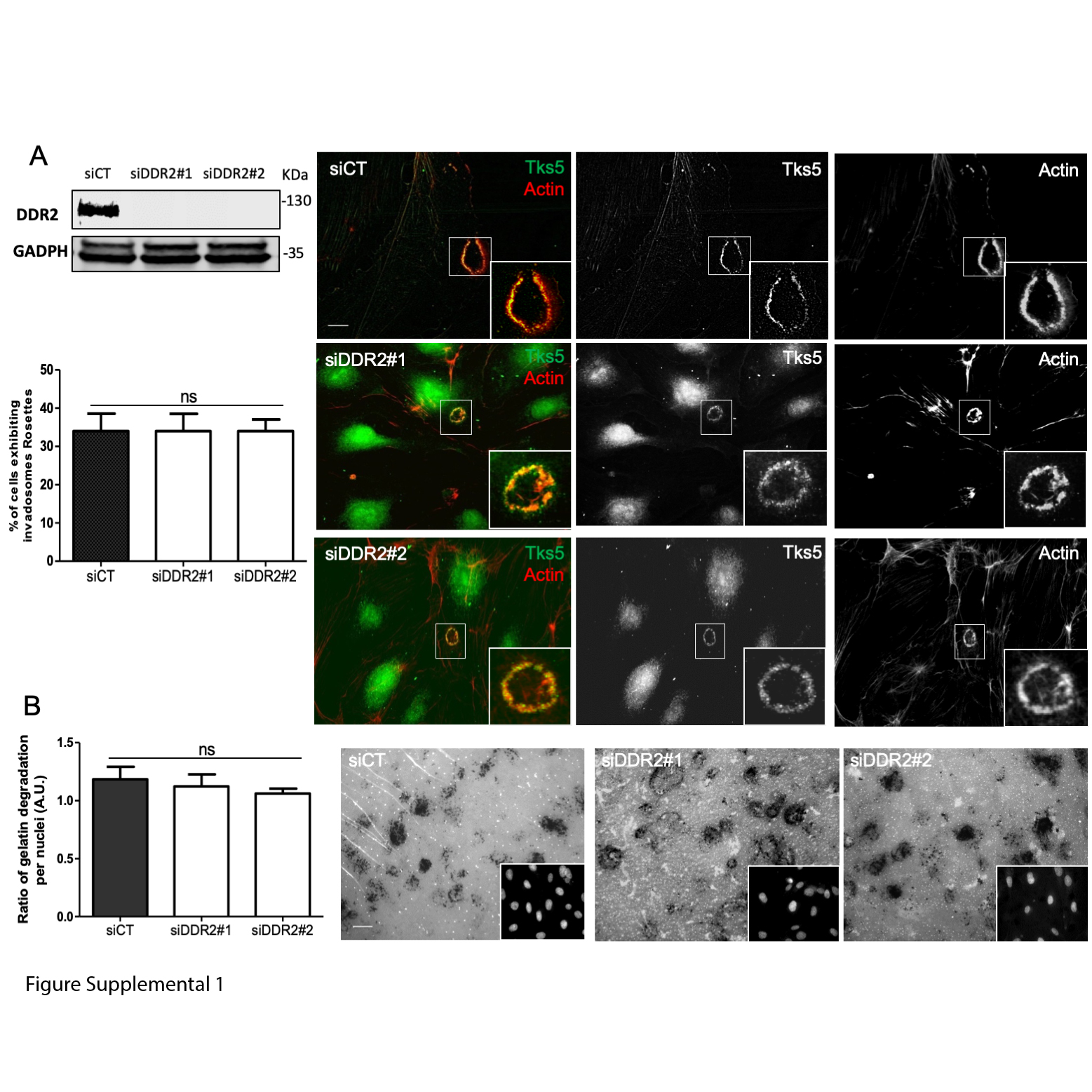

### fig sup 2.jpg

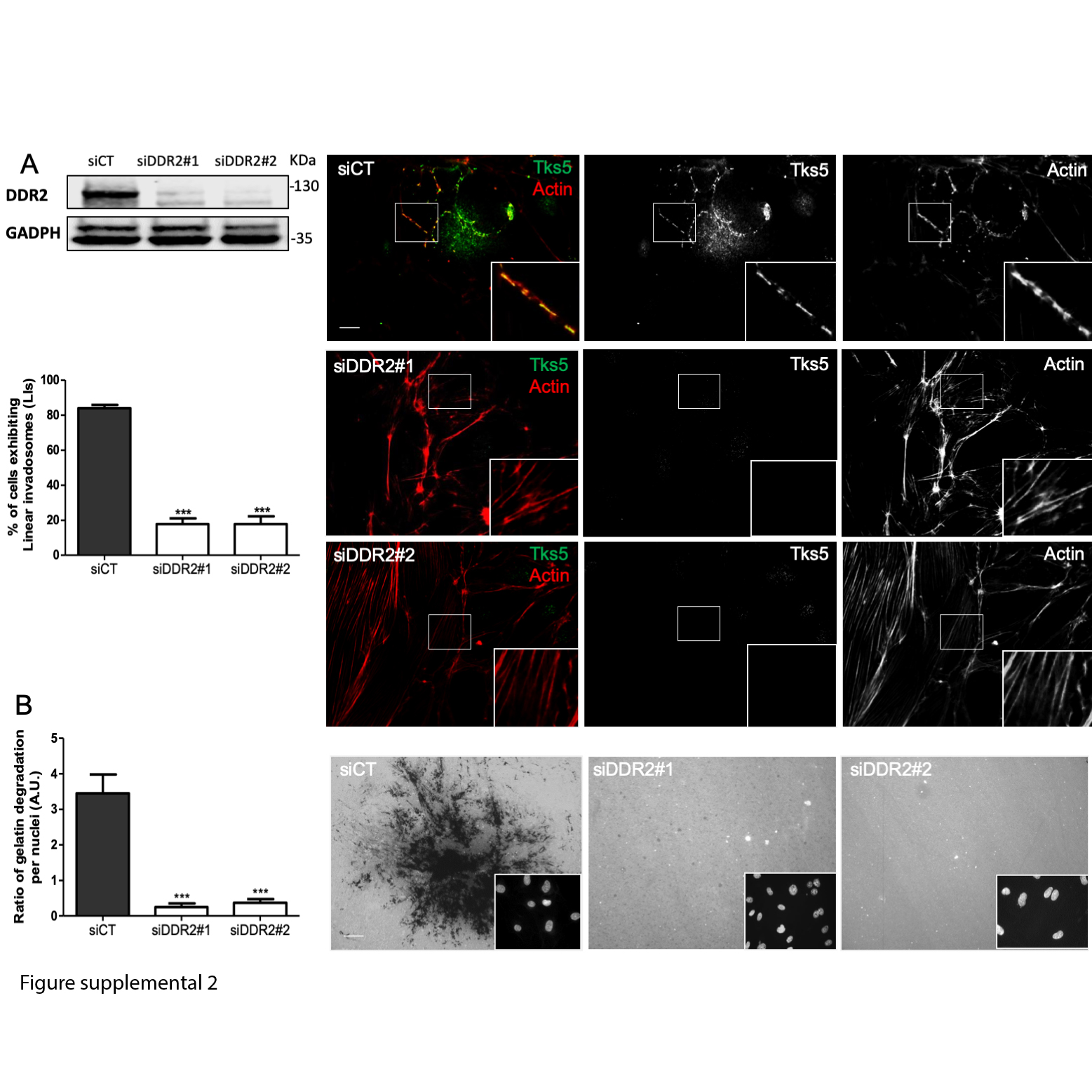

### fig sup 3.jpg

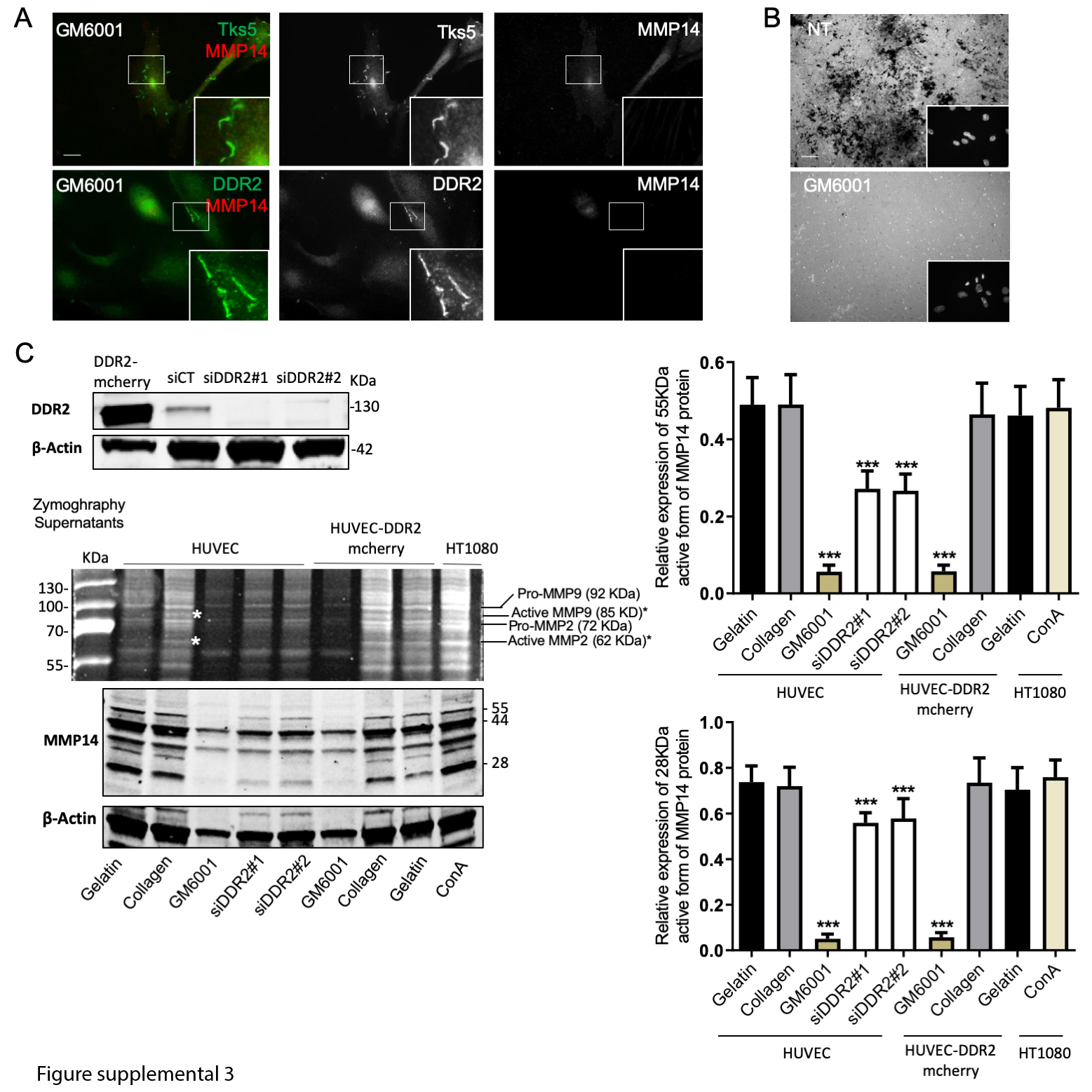
